# Viral surface geometry shapes influenza and coronavirus spike evolution through antibody pressure

**DOI:** 10.1101/2020.10.20.347641

**Authors:** Assaf Amitai

## Abstract

The evolution of circulating viruses is shaped by their need to evade antibody response, which mainly targets the glycoprotein (spike). However, not all antigenic sites are targeted equally by antibodies, leading to complex immunodominance patterns. We used 3D computational models to estimate antibody pressure on the seasonal influenza H1N1 and SARS spikes. Analyzing publically available sequences, we show that antibody pressure, through the geometrical organization of spikes on the viral surface, shaped their mutability. Studying the mutability patterns of SARS-CoV-2 and the 2009 H1N1 pandemic spikes, we find that they are not predominantly shaped by antibody pressure. However, for SARS-CoV-2, we find that over time, it acquired mutations at antibody-accessible positions, which could indicate possible escape as define by our model. We offer a geometry-based approach to predict and rank the probability of surface resides of SARS-CoV-2 spike to acquire antibody escaping mutations.

## Introduction

The COVID-19 pandemic, caused by the SARS-CoV-2 coronavirus, is one of the most challenging global health crisis within the last century (1). The virus emerged as a result of a zoonotic shift (2, 3). It is a member of the betacoronaviruses family (4), related to coronaviruses found in Bats (5), and to SARS CoV that cause severe respiratory syndrome (6) as well as other widely circulating members of the family that cause the common cold (7).

Coronaviruses (CoVs) have the largest genomes among RNA viruses (8). Nonstructural protein 14 (nsp14), a subunit of the replicase polyprotein encoded by CoVs is thought to provide a form of proofreading activity that could support the expansion of large CoVs genomes to their current size. One result of such proofreading activity is that CoVs genomes are less mutable compared to other RNA viruses (9), and thus the sequence diversity of SARS-CoV-2 is quite low (10).

In response to the SARS-CoV-2 pandemic, many approaches for antibody (Ab) therapies, and vaccines are being explored (11). Almost all vaccination approaches aim to use the glycoproteins or spike protein (S) of the virus in its trimeric form (12) or vaccinate with the full (inactivated) virus (13). The spike, a class I fusion protein, mediates entry to the host cell by binding to the angiotensin-converting enzyme 2 (ACE2) receptor (Ou et al., 2020) and is the main target of Ab response (Robbiani et al., 2020; Wu et al., 2020). These therapeutic approaches, hopefully, would be able to elicit strong Ab and T cell response against the virus. In particular, Abs against the spike receptor-binding domain (RBD) have been shown to have neutralization and protective capabilities (14, 15).

Since SARS-CoV-2 virus introduction into humans is recent, it probably has not yet evolved extensively to acquire escape mutations from the commutative Ab pressure of the human population (16). One mutation at the spike (D614G) is now widespread and is thought to support a high viral growth rate (17). However, other members of the coronavirus family have been circulating in human populations for many years (18) and evidence of antigenic drift is seen in SARS-CoV-1 (Guan et al., 2003; Song et al., 2005), and among common cold coronaviruses 229E (Chibo and Birch, 2006). Hence, given the prevalence of SARS-CoV-2, to inform vaccine design and understand how the fitness landscape of the virus evolves, it is important to recognize antigenic drift due to Ab pressure if it were to occur. More generally, antigenic drift due to Ab pressure is common in other RNA viruses such as the seasonal influenza virus (19, 20).

Here we sought to understand and predict, from first principle, to what extent the mutability of the spikes of influenza and close relatives of SARS-CoV-2 could be attributed to Ab pressure. The magnitude (titers) of Ab response against a given epitope is a direct consequence of the B immunodominance hierarchy patterns of an immunogen, which are the result of various aspects of the humoral response to antigen (21–23). Amongst them is the B cell repertoire – the number of B cell clones targeting different epitopes (24–29), their germline affinity (24, 30), and T cell help to B cell (31). Here, we concentrate on the geometric presentation of the spike to Abs. We have previously shown using coarse-grained molecular dynamics simulations, that the geometry of the immunogen spike presentation on the virus recapitulates the known immunodominance of hemagglutinin (HA) head compared to its stem (24).

We developed here an in-silico approach to estimate the Ab targeting - a proxy for B cell immunogenicity (24), of residues on the spike surface, and the differential accessibility to antigenic epitopes due to the geometrical presentation of spikes on the surface of the virus. Superimposed on the spike surface, the immunogenicity score gives the *Ab affinity maps* of influenza and corona spikes, which we applied to predict how the antigenic space is explored unevenly across the surface of these glycoproteins. We then used sequences from public repositories (www.ncbi.nlm.nih.gov, www.gisaid.org) to evaluate the *mutability maps* of those glycoproteins. Next, we developed a computational approach based on spectral clustering to compare these maps. We found that about 50% of the mutability maps variability of the S protein of the severe acute respiratory syndrome-related betacoronavirus (sarbecovirus), and 67% of the variability in the mutability of the seasonal influenza spike (HA) can be attributed Ab pressure, as estimated from the model. This suggests average, polyclonal Ab pressure was consequential in the diversification of the coronavirus sarbecovirus spike and the seasonal flu spike. Moreover, our data suggest that the geometry of spike presentation on the viral surface is a major factor determining its mutability.

We further studied the time evolution of SARS-CoV-2 spike mutability up to November 1^st^, 2020. While the overall correlation between our model and the mutability map is still low (correlation coefficient 0.46), we find that it has been gradually increasing over time. While very preliminary, it could suggest that some variants with Ab escape mutations are establishing in the population. Overall, our approach allows us to recognize from first principle, based on the 3D structure of glycoprotein and cryo-EM images of the viral surface, whether their mutational landscape has features suggesting Ab evasion, and rank surface residues according to their likelihood to acquire Ab-escaping mutations in the future. Importantly, this approach can detect early signs of SARS-CoV-2 and influenza adaptation to evade immune pressure by memory B cells.

## Results

### Geometry-dependent affinity of Abs to HA epitopes

The high-density presentation of spikes on the viral surface shelters, through steric impediments, immunologically recessive and conserved residues from Ab targeting – for example, residues belonging to the stem of HA (20, 29, 32). To study the relative accessibility of residues on the spike surface, we employed MD simulations to define how Ab on-rate, and hence affinity, could be modulated by the presentation of the spike. We first studied two geometrically distinct HA-presenting immunogens: 1) Presentation as soluble full-length HA trimer in its closed form [A/New Caledonia/20/1999 (NC99)] (25, 33–35) (Figure 1A-i); 2) HA presentation within H1N1 influenza A (NC99) virus model (Figure 1B-i) [See Material and Methods]. For each presentation form, based on structural considerations, we computed the on-rate for Abs engaging different surface epitopes (Figure 1A-ii, B-ii). The events occurred in the following sequence: 1] a single Ab arm first engages a target epitope; 2] the Ab molecule continues to fluctuate, allowing the second arm to target a second epitope, when the immunogen geometry is favorable, to bind it with a high rate, resulting in bivalent binding (S Figure 2A-B).

**Figure 1.**
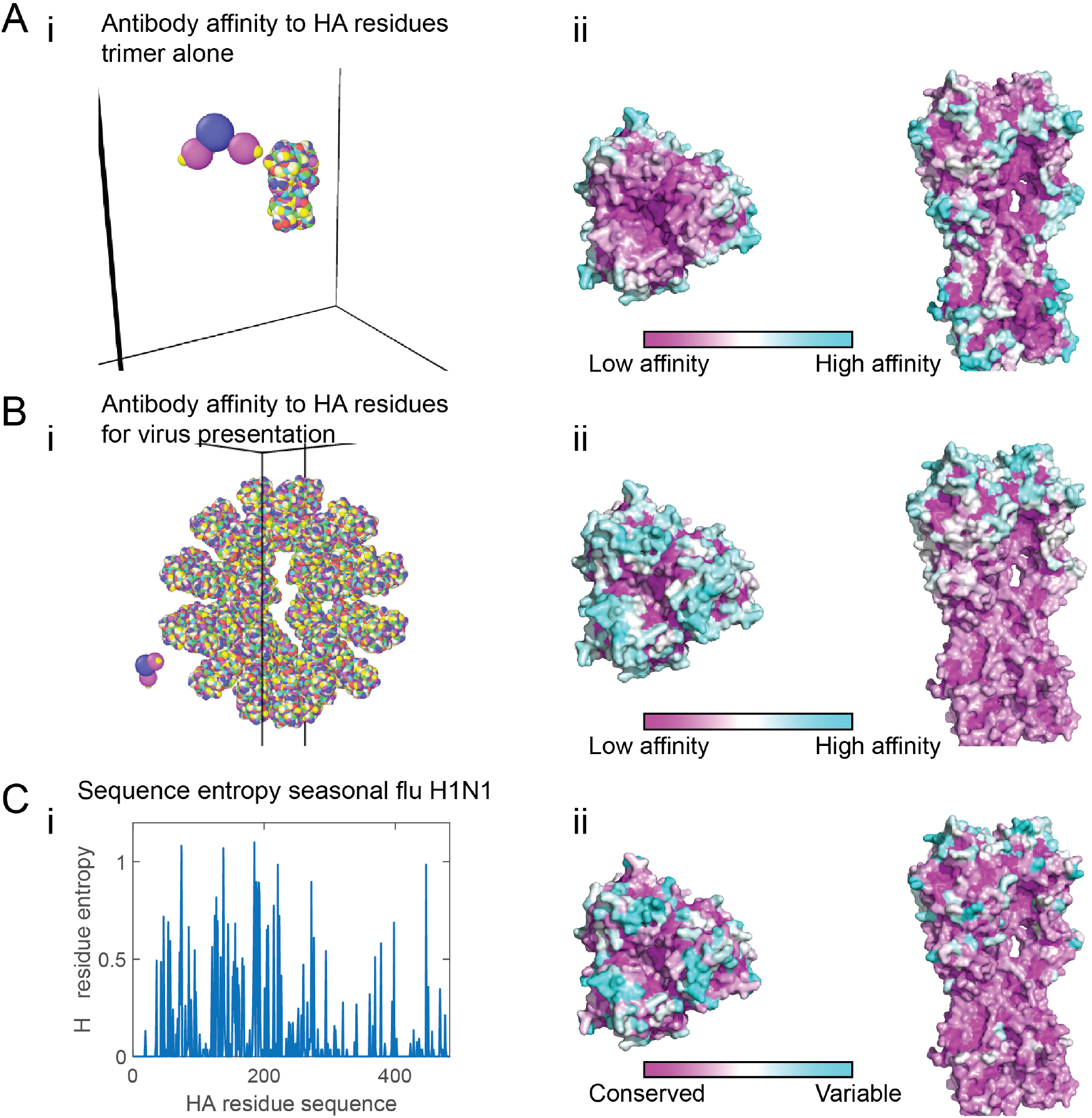
Antibody targeting and mutability of the hemagglutinin protein for the seasonal flu. **(A-B)** Coarse-grain model of the hemagglutinin trimer of A/New Caledonia/20/1999 (NC99) H1N1 influenza protein in its closed form (A). The virus has 40 HA molecules at a spike spacing of 14.8 nm. [Measured spike spacing on influenza is 14 nm (74)] (B). For each immunogen geometry (trimer - A or full virus - B), a detailed atomistic structure of the immunogen is coarse-grained and presented in rainbow colors (panel i). Here every colored bead on the immunogen is a residue, representing a different HA epitope (184 different possible sites on trimeric HA). The antibody structure is presented as the Fc (blue bead), and two antigen binding sites (magenta beads). Panels ii within A-B depict coarse-grained simulations for the Ab affinity (on-rate of Ab first arm binding – see eq.(S4)) to these residues. The affinity estimated from the simulation is superimposed on the HA structure. Top view (left), side view (right). The affinity to cyan sites is high, intermediate to white sites, low for purple sites, and was the average over multiple simulations. **(C)** Panel i depicts the entropy (see eq.(1)) of HA epitopes computed for the seasonal flu (pre-pandemic influenza H1N1 (1918–1957 and 1977–2009) (sequences from (41)). Panel ii shows the entropy of the residues superimposed on HA structure, where highly mutable residues are in cyan, intermediate in white and conserved sites in purple.

We superimposed the in-silico estimated affinity (on-rate of the first Ab arm) on HA structure to represent its *affinity map* (Figure 1A-i, B-i). In the context of the free HA trimer presentation, we found that residues at convex sections on the spike surface were more accessible to the Ab, resulting in high affinity (Figure 1A-ii). In the context of virus HA presentation (Figure 1B-ii), similar behavior followed, and the density of spikes on the viral surface reduced the ability of the Ab to penetrate and interact with epitopes on the lower part of the spike, resulting in an affinity gradient of the Abs targeting residue along the main axis (Figure 1B-ii right). Hence, presentation on the virus surface, as occurs in vivo, leads to an immunodominance or Ab pressure (targeting) gradient along the main axis of HA.

### Antibody pressure directs the evolution of the seasonal flu

Viral infection elicits a humoral response and the production of Abs that target residues on the surface of glycoproteins. For circulating viruses to propagate in a population, they have to evade neutralization and recognition by Abs (36, 37). To do so, they accumulate mutations on their glycoproteins (38, 39). Because sterically hidden residues are less accessible to Abs, we hypothesized their need to mutate is smaller compared to more accessible ones. Hence, we expected spike evolution and the mutational landscape to follow Ab pressure.

The influenza virus mutates from one year and the next, where most of the mutations are concentrated in five antigenic sites (Sa, Sb, Ca1, Ca2, Cb) located at the head of HA (40). Along with genetic drift, escape from neutralization by Abs is one of the main factors contributing to HA mutability (19, 20). To examine the relationship between Ab pressure as defined by our model, and HA surface mutability, we studied the evolution of the seasonal influenza virus H1N1 using sequences dating back to 1918 (see Material and Methods). Following alignment, we computed the entropy of each surface residue identified as an epitope (see Material and Methods). The entropy *H_j_* of residue *j* is given by

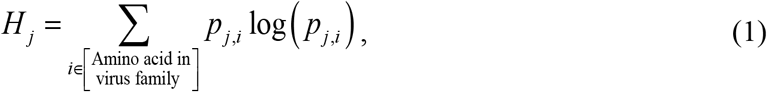

where *p_j,i_* is the probability of amino acid *i* to appear at residue *j* across the viral population (Figure 1C-i). By superimposing the residue entropy on the surface of HA, we created its *mutability map* (Figure 1C-ii). Interestingly, the mutability map is comparable to the affinity map computed for the virus presentation (Figure 1B-ii), showing a pattern of diminishing mutability gradient along the main axis of the spike, but less so to the results for the trimer presentation (Figure 1A-ii). This is corroborated by previous studies showing that the HA head acquires more mutation and evolves faster than its lower part – the stem (41).

We next quantified the similarity of the affinity and mutability maps. Because of the coarse-grained nature of our antibody model, we decided to aggregate close-by residues on the spike surface (Figure 2A). As the surface features of the spike appeared to be an important factor in determining Ab affinity, we applied a non-linear mapping (manifold learning) algorithm - diffusion maps (42) on the epitopes’ positions and used the first three components (Figure 2B). We then applied the k-means clustering algorithm (43) (spectral clustering) to aggregate residues in this space into epitope clusters (Figure 2C). We computed for each epitope cluster *k* its entropy and affinity as follows:

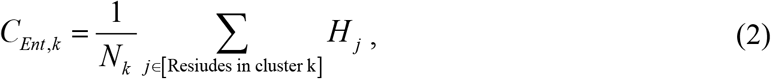

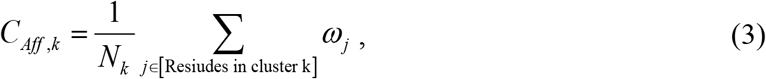

where *C_Ent,k_* and *C_Aff,k_* are the epitope cluster entropy and affinity respectively, *N_k_* is the number of residues in cluster *k*, *ω_j_* is the affinity (on-rate of the first Ab arm to epitope) to residue *j*, and the entropy *H_j_* of residue *j* is given by eq.(1).

**Figure 2.**
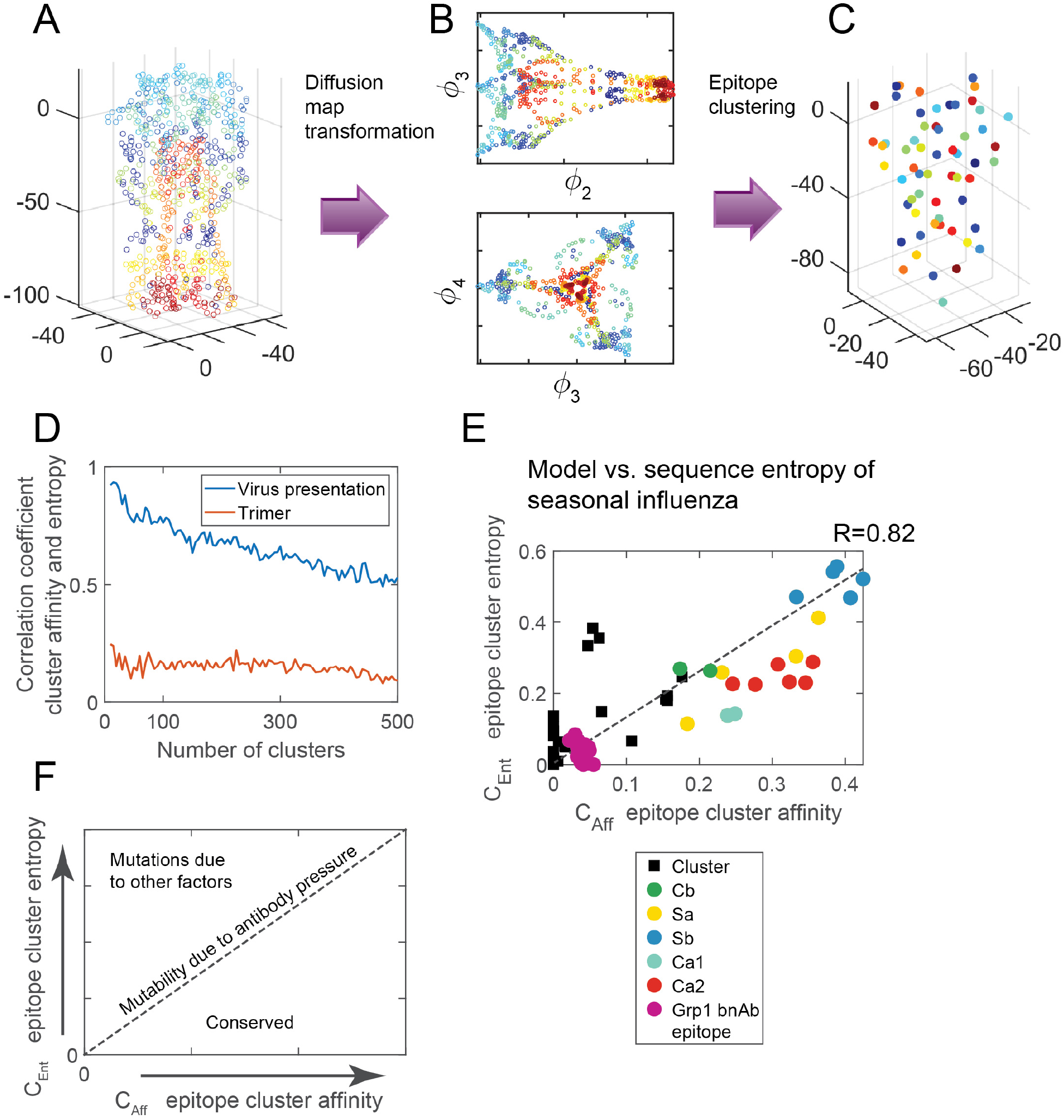
Antibody pressure guided the mutability of the hemagglutinin. **(A)** HA protein. Each circle corresponds to a surface residue (epitope) and has a different color. **(B)** 2d projections of the first four eigenvectors of the epitope positions following diffusion map decompositions. **(C)** Surface residues (epitopes) were clustered using k-means clustering algorithm applied to spectral decomposition shown in B. Same color-coding as in (A) **(D)** The correlation coefficient between epitope cluster entropy (eq.(2)) and the epitope cluster affinity (eq.(3)), as a function of cluster number, computed for HA in the virus presentation depicted in Figure 1B (blue), and at the trimer presentation depicted in Figure 1A (red). **(E)** Scatter plot of the epitope clusters entropy computed for the seasonal influenza H1N1 vs. the epitope clusters affinity. The correlation coefficient between them is 0.82. Marked are clusters containing residues belonging to the five known antigenic sites of flu (Cb - green, Sa - yellow, Sb - blue, Ca1 - cyan, Ca2 - red). Also marked is the group 1 conserved broadly neutralizing antibodies epitope (purple). (The number of clusters is 60). **(F)** Schematic of the relationship between entropy and computed Ab affinity for circulating viruses evolving under Ab pressure.

To assess the predictive strength of the computed affinity in explaining the observed mutability, we computed the correlation between *C_Aff,k_* and *C_Ent,k_* as a function of the cluster number (Figure 2D). We find that the correlation values for HA virus presentation are high, with a maximum of 0.92 for 10 clusters, suggesting the in-silico model can explain a significant fraction of the mutability. Interestingly, the correlation value, regardless of cluster number, is always larger for the virus presentation than for the trimer (Figure 2D), highlighting that spike evolution and escape due to Ab pressure occurs in the context of the virus - as a result, both mutability and Ab affinity vary most along the main axis of the spike.

To determine the optimal number of clusters *k* for comparison between the two maps, we first estimated the Total Within Sum of Squares for different cluster numbers (S Figure 5A) and used the elbow method to choose *k* = 60 (44). For *k* = 60, we found a correlation of 0.82 between *C_Ent,k_* and *C_Aff,k_*, suggesting that epitope cluster affinity, at this resolution, can explain 67% of the variability in the mutability map of HA.

Surprisingly, most epitope clusters that contain residues belonging to the five antigenic sites show a linear relation between their entropy and affinity, suggesting that the mutability of these sites simply follows from their position on HA, the geometry of its presentation on the viral surface, and is due to Ab pressure (Figure 2E). Epitope clusters containing conserved residues at the HA stem belonging to the HA Group 1 broadly neutralizing epitope (20, 29, 32) similarly align.

Taken together, these results suggest that the mutability of surface spike epitopes of circulating viruses can be described using a phase diagram (Figure 2F). The mutability of epitope clusters that lay on a linear line of epitope cluster entropy vs. epitope cluster affinity is related to the average Ab pressure acting on these residues (Figure 2F). Epitope clusters below the line are more conserved than would be expected based on their accessibility to Ab pressure and could be due to the presence of functionally important sites. Epitope clusters above that line are more mutable than would be expected due to Ab pressure and may result from allosteric immune escape (45), escapes from CD8+ T cells (46, 47), glycosylation (48), or other factors.

### The mutability map of the sarbecovirus spike follows geometry-dependent antibody pressure

Coronaviruses are capable of crossing the species barriers in zoonotic shifts resulting in the SARS-CoV pandemic in the year 2002-2004 and the 2012 MERS pandemic (4). To understand whether the geometrical principles controlling the distribution of mutation on the spike surface are general across species, we applied our computational model to study the mutability of the spike protein of close relatives of SARS-CoV-2 – the sarbecovirus subgenus. We considered two presentations of the corona spike (S protein) to Abs: 1) Presentation as soluble full-length S trimer in its closed form (49) (S Figure 3A-i); 2) S presentation on the coronavirus surface (Figure 3A-i) [based on the cryo-EM structure of SARS that has 65 spikes on its surface (50) and SARS-CoV2 spike (49) (See Material and Methods)].

**Figure 3.**
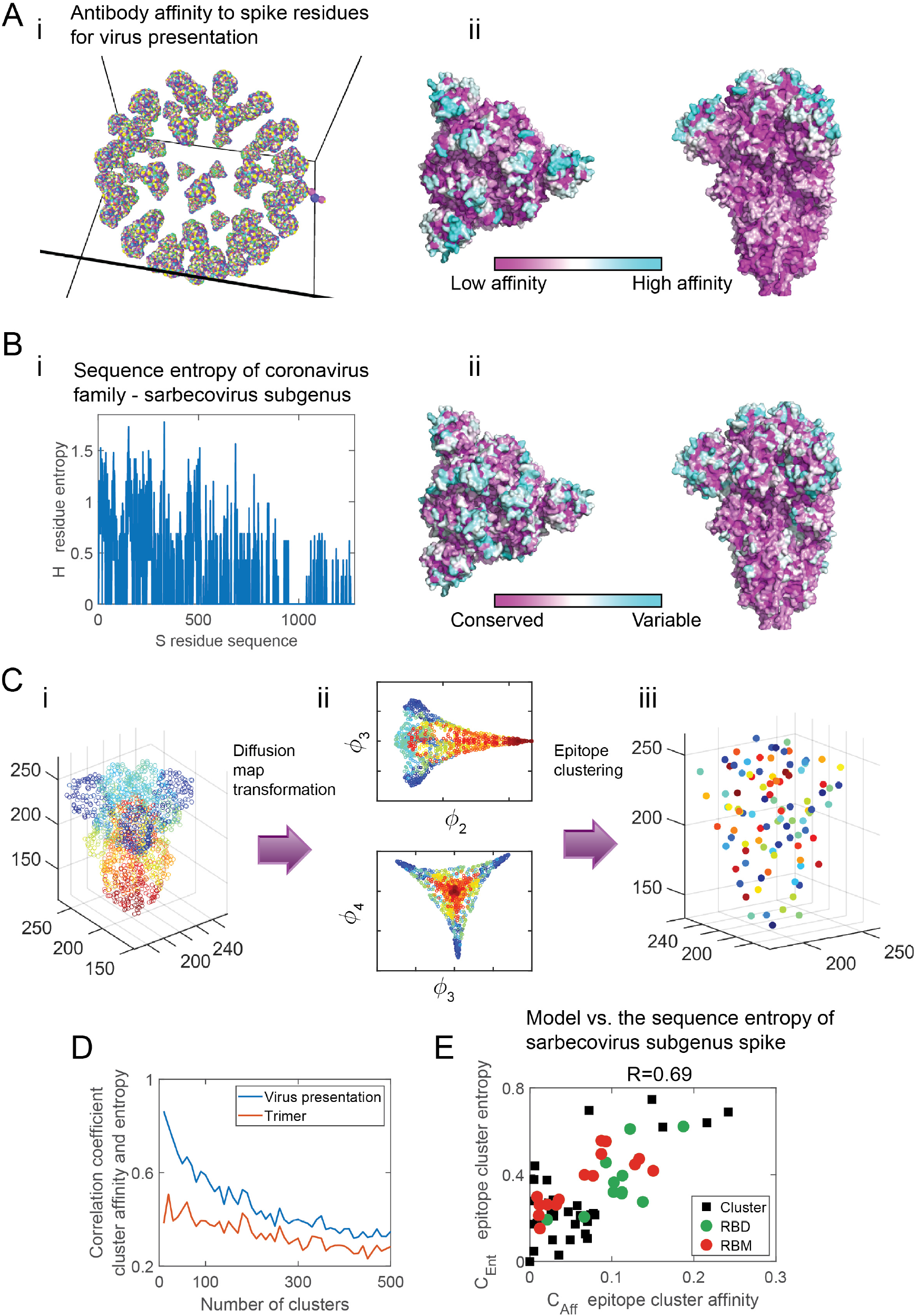
Antibody targeting and mutability of the sarbecovirus subgenus spike. **(A)** SARS-CoV-2 spike (S) protein in its closed form (49) arranged on a sphere to create a virus model. The virus model has 65 S molecules at a density of 0.27 spikes per 100nm^2^ (50). A detailed atomistic structure of the spike is coarse-grained and presented in rainbow colors (panel i). Every colored bead on the spike is a residue, representing a different S epitope (254 different possible sites on trimeric S). Panel ii depicts coarse-grained simulations for the Ab affinity to these residues (see Figure 1A-ii for definition and color-coding). **(B)** Panel i depicts the entropy (see eq.(1)) of each spike residue computed for the sarbecovirus subgenus spike (see Table 1). Panel ii shows the entropy of the residues superimposed on the spike structure. Same color-coding as in Figure 1C-ii. **(C)** Pane i. The spike protein of the coronavirus. Each circle corresponds to a surface residue (epitope) and has a different color. Panel ii. 2d projections of the first four eigenvectors of the epitope positions following diffusion map decompositions. Pane iii. Clustering of the surface residues of the spike protein using k-means clustering algorithm applied to the spectral decomposition shown in panel ii (*k* = 60). **(D)** The correlation coefficient between epitope cluster entropy (eq.(2)) and the epitope cluster affinity (eq.(3)), as a function of cluster number, computed for the corona spike in the virus presentation (blue), and at the trimer presentation (red). (see S Figure 3 for the trimer presentation affinity map) **(E)** Scatter plot of the epitope clusters entropy, computed for the sarbecovirus spike vs. the epitope cluster affinity estimated from the simulations. The correlation coefficient between them is 0.69. Clusters that contain residues belonging to the RBD are in green and those containing residues belonging to the RBM are in red. (The number of clusters is 60).

We first used in-silico simulation to estimate the Ab affinity to surface residues of S, when presented as a trimer or the surface of the virus model (See Material and Methods). Similarly to our observation for the Abs affinity against HA, we found an increased affinity to convex regions, an affinity gradient along the main axis of S for the virus presentation (Figure 3A-ii), but not for the trimer presentation (S Figure 3A-ii). Next, we analyzed sequences of close relatives of the SARS-CoV-2 spike within the sarbecovirus subgenus (Table 1). Following alignment and construction of the phylogenetic tree (S Figure 4), we computed the mutational entropy of each surface residue identified as an epitope using eq.(1) (Figure 3B-i) and superimposed it on the spike surface to create its mutability map (Figure 3B-ii). We observed that the most significant change in mutability is along the main axis of S. To quantitatively compare the affinity and mutability maps, we applied the diffusion map transformation on S and clustered the epitopes (Figure 3C). Studying the correlation value as a function of cluster size (Figure 3D), we found that the correlation between the model and the mutability map is always higher for the virus spike presentation compared to the trimer, highlighting that the geometrical context in which Abs interact with the spike determines its mutability. We found a high degree of correlation (R=0.69) between *C_Ent,k_* and *C_Aff,k_*, suggesting that affinity as computed by our model, at this resolution, can explain 48% of the variability in the mutability map of S (Figure 3E). The high degree of correlation between entropy and affinity suggests that average Ab pressure shaped, to the first order, the mutability of the sarbecovirus subgenus spike. While for seasonal influenza, the HA entropy (Figure 1C) was the result of a gradual accumulation of mutations over time, S protein entropy (Figure 3B) analyzed here is the result of a horizontal mutational process occurring simultaneously in different hosts, suggesting the virus evolves under similar geometrical immunoglobulin pressure.

**Table 1.**
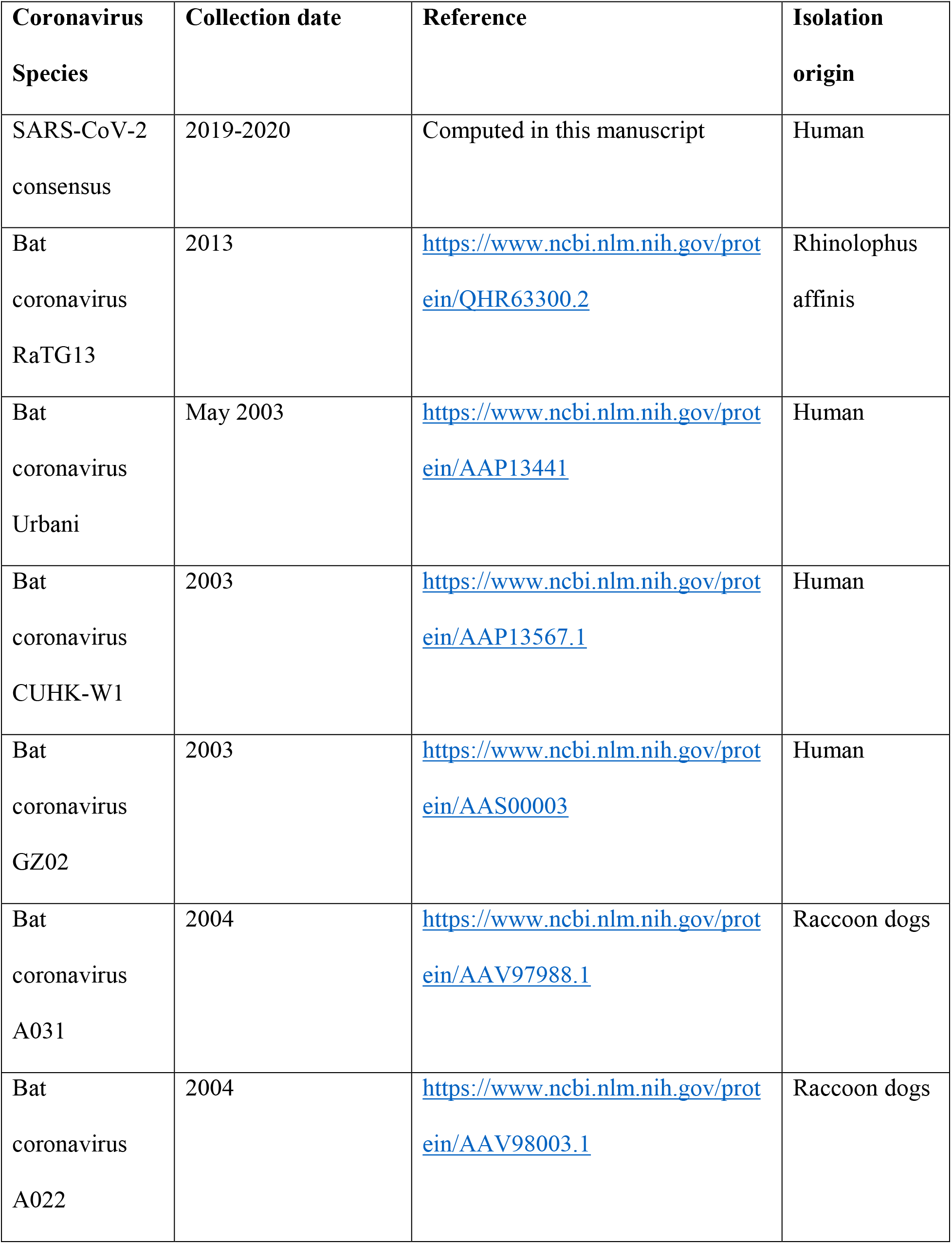
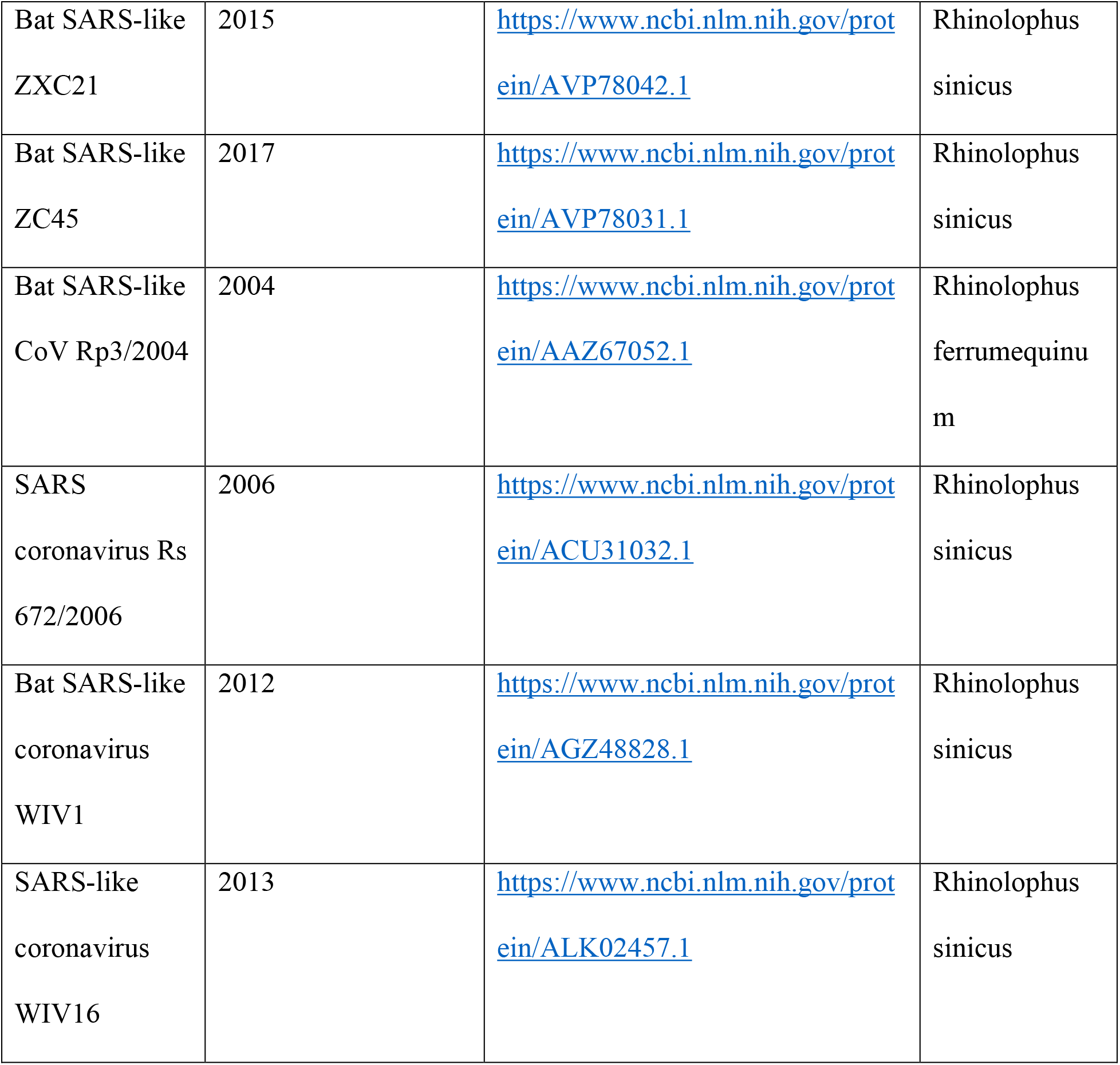
Sarbecovirus. Species used for the analysis detailed in Figure 1C.

The receptor-binding domain (RBD) is involved in the spike binding to ACE-2 (51, 52). It has been shown that neutralizing Abs targeting the RBD can offer protection (14). Within the RBD, residues belonging to the receptor-binding motif (RBM) are most important in binding to ACE-2. We recognized epitope clusters to which residues part of the RBD and RBM belong (Figure 3E). Many of the epitope clusters have both high entropy and high affinity, which could suggest mutations acquired at these key domains across the spike are due to evasion from Abs, as well as adaptation to the host-specific receptor. Several of the highly targeted and mutable epitope clusters are not part of the RBD. Hence, Abs targeting these residues will not necessarily offer neutralization activity. However, Abs targeting these clusters can control viral infection through non-neutralizing pathways (53), thereby motivating the virus to mutate these highly targeted parts.

### SARS-CoV2 and the 2009 influenza pandemic spikes mutability is not predominantly due to antibody-pressure

Our analysis suggested that spike presentation geometry is an important factor governing the mutational entropy of viruses circulating either over long periods (influenza) or across species (sarbecoviruses). To find whether this observation can be generalized to pandemics, we computed the sequence entropy of HA for the 2009 flu pandemic H1N1 (sequences from (41), GISAID) (Figure 4C-i). Superimposing the entropy on the HA structure (Figure 4A-ii), we did not observe immunodominance gradient along the main axis of HA observed in the computational model (Figure 1A-ii). Unlike for the mutability of seasonal flu, the correlation coefficient between epitope cluster pandemic entropy and epitope cluster affinity was low (0.18) (Figure 4A-iii).

**Figure 4.**
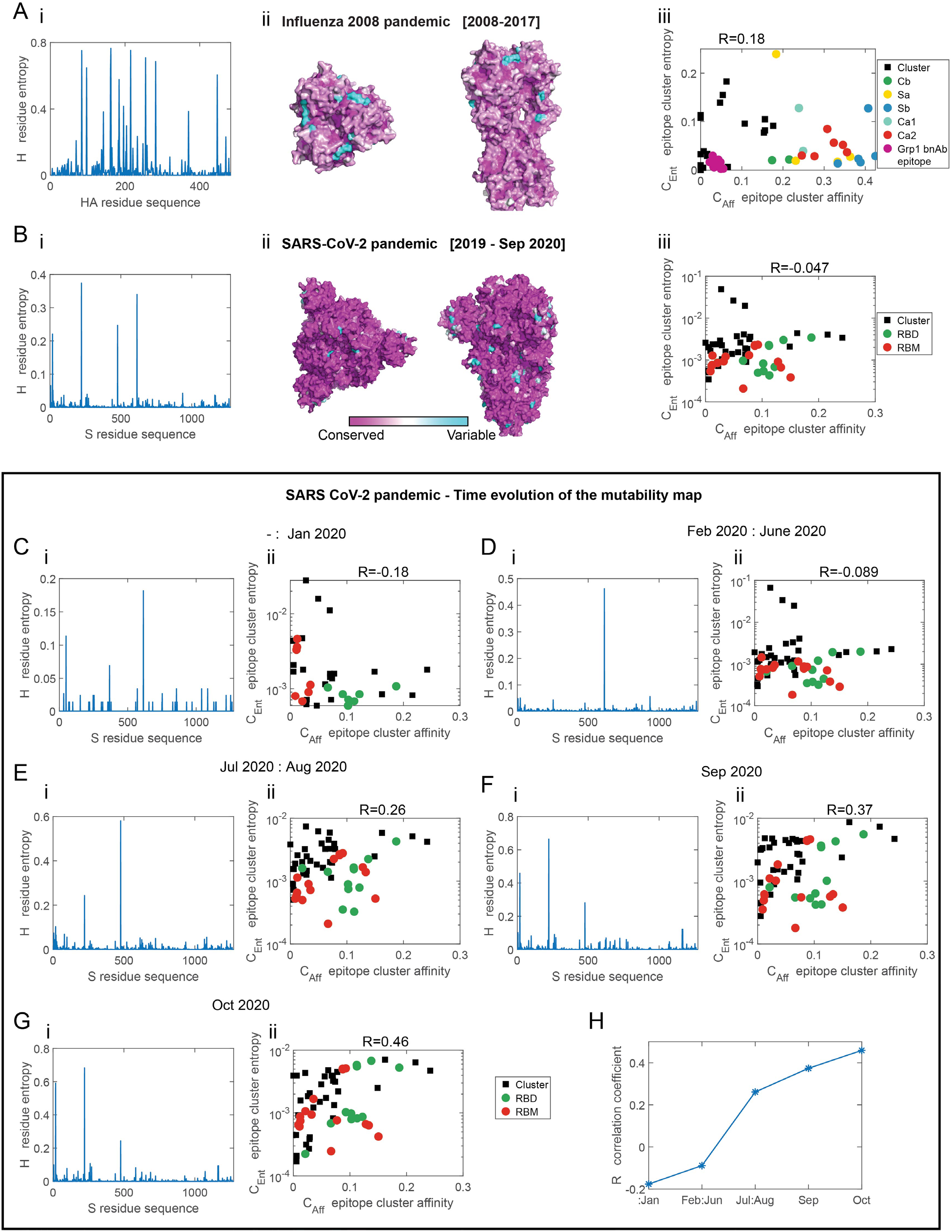
Spike evolution of the 2009 influenza pandemic and SARS-CoV-2. Comparison of the mutational variance of sequences (mutability map) and the affinity maps computed from the computational model. For A-B, panel i depicts the residue entropy as a function at different positions. For A-B, panel ii depicts the entropy of the residues is superimposed on the spike. Same color coding as in Figure 1C-ii. Panel iii. Scatter plot of the entropy of epitope clusters, against the epitope cluster affinity computed for the spike. **(A)** Sequence entropy of HA for the pandemic flu H1N1 (2009–2017) (sequences taken from www.gisaid.org, and (41)). The correlation coefficient between the epitope cluster entropy and affinity is R=0.18. **(B)** (i) Sequence entropy of the S spike protein of SARS-CoV-2 computed for all S protein sequences up-to October 31^th^ 2020 (sequences downloaded from www.gisaid.org). (ii) The correlation coefficient between the epitope cluster entropy and epitope cluster affinity is R=-0.076. Same legend as Figure 3E. **Time-dependence of SARS-CoV-2 sequence entropy.** The entropy of the S spike protein of SARS-CoV-2 computed for sequences collected at 5 time periods since the beginning of the pandemic (panel i) and correlation to the computationally derived affinity map, following epitope clustering (panel ii) (same clusters as those shown and used in Figure 3C-iii, E). **(C)** up to February 1st 2020, R=-0.18, **(D)** February-June 2020, R=-0.089 **(E)** July-August 2020, R=0.26, **(F)** September 2020, R=0.37, **(G)** October 2020, R=0.46. **(H)** The correlation coefficient as a function of time. (Find an interactive, comparison of the time-dependent mutability map to the affinity map here https://amitaiassaf.github.io/SpikeGeometry/SARSCoV2EvoT.html).

SARS-CoV-2 zoonotically shifted to humans in 2019 (5), probably from Bats via pangolins, although its precise evolutionary path is still unclear. Since then, it has spread in the human population, infecting more than 35 million people as of October 2020. Because of its proofreading capability, the virus evolution is slow. To analyze its total mutational entropy up to November 1^st^, 2020, we downloaded publically available SARS-CoV-2 sequences from GISAID (www.gisaid.org) (54) (sequences choice is discussed in Material and Methods), computed the sequence entropy (Figure 4B-i), and superimposed it on the close structure of the spike (Figure 4B-ii). Interestingly, the mutability map does not show the same gradient pattern as observed for the sarbecovirus subgenus spike entropy (Figure 3B-ii). We applied the same clustering (*k* = 60) to compare the epitope cluster entropy and affinity and found a low value of the correlation coefficient (−0.047) (Figure 4B). Hence, the total sequence entropy of SARS-CoV-2 thus far is not dominanted by escape from Ab mutations.

### Time evolution of SARS-CoV-2 mutability map

To see if we could observe changes in the evolution trend of the virus of the time, indicative of Ab escape mutations, we separated SARS-CoV-2 sequences into five groups based on the time at which they were captured: 1] before 02/2020, 2] 02/2020-06/2020, 3] 07/2020-08/2020, 4] 09/2020, and 5] 10/2020 (Figure 4C-G). Computing the correlation coefficient between the Ab affinity map and the sequence entropy, we found a significant increase over time (Figure 4H) from a value of −0.18 in Feb 2020 to 0.46 for sequences sampled during 10/2020. While the correlation value of 0.46 is still low, this may point that escape from Abs mutations are starting to be a more significant part of its population, in accordance with other reports (16, 55). An increase over months and years in correlation value between the affinity map computed by our model and the evolving mutational landscape of SARS-CoV-2 could indicate that its mutability patterns are being shaped by Ab pressure. (See time dependence here https://amitaiassaf.github.io/SpikeGeometry/SARSCoV2EvoT.html).

## Discussion

Humoral immunity is often characterized by dominant versus recessive responses to different epitopes on the same antigen. This hierarchy of B cell immunodominance depends on many factors, amongst them are the precursor frequency within the germline B cell repertoire, BCR affinity, and the steric accessibility or antigen geometry. Pathogens take advantage of antigen geometry to shield sites of vulnerability. Such is the case in influenza spike hemagglutinin, where conserved sites are located and the sterically hidden stem (56–58), or on HIV spike gp120 where the vulnerable and evolutionary conserved CD4 binding site position does not allow Abs to form bivalent interactions, reducing the effective affinity of Abs (59). While mature Abs are nevertheless capable of approaching sterically restricted sites via somatic hypermutations that could extend, for example, their CDR3 loops (60), immunogen shape and valency manipulates B cell immunodominance patterns, their selection process in the germinal center or the expansion of memory B cell population (24, 61, 62). Because viruses must evade Ab response to survive, B cells immunodominance patterns could be a proxy to glycoprotein mutability, coupled through antigen geometry. Thus, we studied whether spike presentation geometry to Abs is a good predictor of their mutability. Using a coarse-grained model of an antibody, HA, and the S protein of SARS-CoV-2, in both trimer and viral presentation model system, we computed the Ab affinity maps as a proxy for Ab pressure on the spike. We used those maps to assess whether the magnitude of mutability at a cluster of sites is just what would be expected by geometrical considerations (Figure 2F).

We found that for the seasonal flu spike – HA, geometry through the presentation on the virus could explain, to the first order, the mutability patterns at its surface. In particular, the mutability of the five antigenic sites is ordered as would be expected by the geometric restriction imposed by their position on the spike, as did the conserved group 1 epitope, which is functionally important for HA conformation change (Figure 2E). Hence, we speculate that rather than maintaining functionally important sites conserved by negatively selecting mutants at such sites, the virus positions functional sites at a location, where their acceptable mutational rate would be determined by their need to escape from targeting by the average polyclonal Ab response.

To understand whether a similar principle governs the mutability of coronaviruses, we created a similar coarse-grained model of the SARS CoV family. As coronaviruses do not mutate much, we decided to analyze its mutability across the virus sub-species, using sequences isolated from different hosts in the years 2003-2019. In mammalians, these viruses have to evade immunoglobulin response which we hypothesized would lead to geometrically similar escape patterns. We found that geometry, through Ab targeting, shapes to the first order the mutability patterns on the sarbecovirus subgenus spike map. Hence, these viruses evolve across various hosts under roughly geometrically-similar Ab pressure – at least the main axis of the virus seems to be the first, principles axis of mutability resulting from the density of spikes on the viral surface.

The mutational probability distribution we sampled for the sarbecovirus subgenus is analogous to sampling different “realizations” of the statistical ensemble of the sequence landscape of the viruses (63), where each realization is a viral from a different host. For the seasonal flu, we considered sequences over a large period - starting from 1918 and aggregate them to a single probability distribution analyzed. In both cases, presentation geometry roughly explained sequence entropy. Comparing both these approaches to describe mutability distribution is conceptually similar to the ergodic theorem in statistical physics, where the averages of a stochastic process sampled over time are equivalent to the averages computed over different statistical realizations. While evolution patterns of mutating viruses are not an ergodic system in general – as many mutants are not viable, and hence unreachable in the sequence space, the similar geometry of immunoglobulins and spike presentations could be is the reason our model works for both these different instances, with mutations distributed across time (for influenza), or across species (for sarbecoviruses). Statistical physics models have been previously used (64) to analyze the sequence space to compute the fitness landscape space of viruses (65, 66). The overall fitness of viruses is often split into its intrinsic fitness of the virus and fitness component related to evasion from the immune response (i.e. Abs) (67). As our approach allows for rough estimation, from first principle, of the virus Ab-dependent element of the fitness, it can be used as a prior in inference methods of the intrinsic fitness.

Because of its proofreading mechanism, SARS-CoV-2 is not expected to mutate much. Nevertheless, since the SARS-CoV-2 pandemic has erupted, its sequences have been analyzed to detect mutations that would increase its fitness, infection capabilities, or allow it to escape from Abs (16, 17, 55, 68). To see if we can find traces of escape due to Ab pressure on the SARS-CoV-2 spike, we compared its mutability map to our computed affinity map over time and found an increase in the correlation value since the beginning of the pandemic (Figure 4H). While the correlation value of 0.46 computed for October 2020 is still low, it could suggest the spike starts to acquire some escape from Ab mutations. Since the RBD is involved in binding to ACE2, mutations at this domain could also modify the binding energy to the receptor. Interestingly, for the 2009 influenza pandemic, we found a low correlation value of 0.18. This could suggest that the 2009 flu pandemic has not been evolving for long enough under Ab pressure for the geometric pattern to be apparent and that the evolution of pandemic viruses is not, at least initially, directed by Ab pressure acting on their surface residues. It is more likely that mutations that accumulate in their spikes serve to increase their fitness and infecting capabilities in humans. Additionally, it is possible that pandemics do not elicit as strong a memory recall as seasonal/circulating viruses, and hence do not need to evolve as rapidly to escape Ab immune pressure.

We propose here a simple geometrical interpretation of the surface mutational landscape of that spike that could inform, based on sequences and the 3D structure alone, whether a dominant component of virus evolution is evasion from Abs. This technique could serve as an indicator of the evolutionary stage in the infection trajectory of a virus and whether it is on its way to becoming a circulating virus such as the seasonal flu.

## Materials and Methods

### The geometry of immunogens and epitope choice

The first input to our model was an atomistic description of the geometry of our immunogens, which we generated from available structural information and pdb files (49, 69). For HA and S, solvent-accessible residues were identified using pymol script “findSurfaceResidues” (https://pymolwiki.org/index.php/FindSurfaceResidues), which identifies atoms with a solvent accessible area greater than or equal to 20 Ang^2^ (HA) and 15 Ang^2^ (S). We then find the residues to which those atoms belong to. This selection criterion gives a uniformly distributed set of residues on the face of HA and S (see S Figure 1A, B). A total of 184 epitopes (residues) were chosen for HA and 254 epitopes for S.

We constructed a simplified model of the influenza virus, in which 40 HA molecules are arranged in a fixed conformation on a sphere of radius equal to 16nm (a value chosen for computational tractability). The model recapitulates (24) the average spacing between adjacent HA on the influenza viral surface of ~ 14 nm (Harris et al., 2013). We also constructed a simplified model of the coronavirus based on the cryo-EM images of the SARS virus, in which 65 S molecules in closed form (49) are arranged in a fixed conformation on a sphere of radius equal to 87nm (50), resulting in a density of 0.27 spike pre 100nm^2^.

Steric constraints affect the access of antibodies to epitopes and this modifies the on-rate, thus modulating the affinity. To compute the relative magnitude of this effect for different epitopes presented by immunogens with different geometries, we employed MD simulations. In these simulations, a Lennard-Jones potential describes the interactions of antibodies with the immunogen atoms, and a separate Morse-potential is used to model interactions of the antigen binding region of the Ab to its specific cognate epitope (see Table S 1). To estimate the steric effects alone, we first assumed that the affinity of Abs to all epitopes was equal in the absence of steric constraints. We then used MD simulations (Lammps software) (70) to compute the average time for the Ab antigen-binding region (S Figure 1C-D) to find the target epitope for the first time, which is called a “first passage time”. By running simulations multiple times, and then averaging over many simulations, we could estimate the mean first passage time to the epitope. The inverse of the mean first-passage time is the on-rate, and thus we computed the relative on-rates for Ab binding to different epitopes for different immunogen geometries. We take the on-rate of the first arm of the Ab model as a proxy for Ab affinity to a residue.

### Coarse-grained model of the antibody

To estimate the encounter probability and rate of different residues on the surface on the immunogens by the Ab, we employed coarse-grained MD simulations. The B cell receptor is represented using 8 beads (see S Figure 1). We used a coarse-grained model of the Ag and Ab (see S Figure 1). In (71), a model of the Ab was suggested, built from ellipsoids and spheres. Here, we built our Ab model using spheres of different sizes to approximate the same dimension and flexibility of the Ab. The MD simulation system is composed of different beads (see Table S 1). This size of the beads was chosen such that the distance between the two Fabs is approximately 15nm and the length of the Ab arm is 7nm (72). The size of the Fc region is chosen to be 5nm (73) (see Table S 1). To construct the 7nm arm we use 3 beads (types 4,5,6 – S Figure 1B-C, Table S 1), where nearest-neighbor beads are connected with rigid bonds of length 1.75nm. Bead type 4 (arm hinge) is connected to bead 3 (Fc hinge) by a rigid bond of length 1.75nm. The epitope bead (type 7, Table S 1) was chosen to have the same size as the Fab beads (1.75nm) (Table S 1). The beads along the arm (type 4,5,6) are on a straight line (no kink), and the middle bead (type 6) is larger, to approximate the elongated ellipsoid shaped arm of the Ab (71).

The average angle between the two arms of the Ab fluctuate with a mean of 120 degrees and obeys the harmonic potential

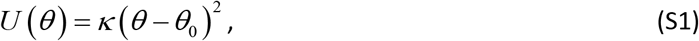

with *θ*_0_ = 0.66radians and *κ* = 10k_b_T /radian^2^, resulting in a relatively rigid model of the Ab (De Michele et al., 2016).

The system is integrated using a Langevin thermostat under “fix nve” to perform performs Brownian dynamics simulations (see https://lammps.sandia.gov/doc/fix_langevin.html).

The Fab bead interacts with the respective epitope bead via the Morse potential

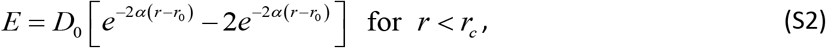

where *r*_0_ = 1.75nm which is the distance between the Fab bead and an epitope bead at which the LJ energy between them is zero, and the cutoff radius *r_c_* = 2.2nm. *D*_0_ = 50 is the energy and the bond fluctuation scale *a =* 1nm^−1^: the Morse potential only serves to anchor the 1^st^ arm to the epitope allowing the second arm to search for a second epitope.

The beads interact with the Lennard-Jones potential

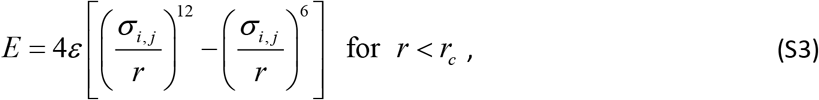

where *ε* = 1, *σ_i,j_* is the interaction distance between beads i and j, and the cutoff radius is *r_c_* = 2^1/6^*σ_i,j_*. The values of *σ_i,j_* are detailed in Table S 2. The LJ interaction distance *σ_i,j_* between all beads composing the Ab arm (types 4, 5, 6), and the epitope bead (type 7) is 1.75nm to construct the 7nm long arm. The LJ self-interaction distance of the Ab arm bead (type 6) was taken to be 4.2nm (Table S 1) to maintain an angle of approximately 120 degrees between the arms. The interaction distance of other pairs of beads is the sum of their radii (Table S 2).

### Estimating the on-rates to the epitopes

The on-rate to each of the residues is estimated using MD simulations. Each simulation runs for a predetermined amount of time and we find the diffusion-limited first passage time of one of the Fabs to the neighborhood of the target residue. The on-rate for the first arm to find an epitope is given by

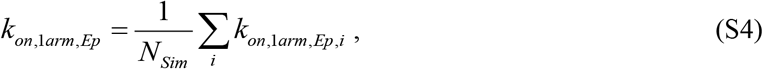

where *k_on,1arm,Ep,i_* is the on-rate estimated from simulation, *i*, for the Ab to find epitope *Ep*, and *N_Sim_* is the number of independent simulations we perform. In a simulation where the Ab does not find its respective epitope we take *k_on,1arm,Ep,i_* = 0. We perform independent MD simulations to estimate *k_on,1arm,Ep_* for each epitope (7 independent simulations for the HA trimer, 12 for the influenza virus model, 17 independent simulations for the S protein trimer, 9 for the coronavirus model).

The on-rate for the binding of the second arm is given by

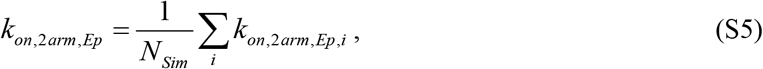

where in a simulation where the Ab does not find its epitope we take *k_on,2arm,Ep,i_* = 0.

### Viral sequences

The sequences analyzed here of the seasonal influenza H1N1 and the 2009 influenza pandemic are from (Kirkpatrick et al., 2018) - which originate from www.gisaid.org. SARS-CoV-2 sequences were downloaded on November 8^th^ 2020 from www.gisaid.org. Out of a total of 189550, only high quality (complete) sequences of length 1274 amino acid (135216 sequences) were analyzed. The consensus sequence of SARS-CoV-2 was calculated using the BLOSUM50 scoring matrix in Matlab. Sequences of the sarbecovirus subgenus were downloaded from www.ncbi.nlm.nih.gov. (see Table 1). The alignment of those sequences was done using ‘GONNET’ scoring matrix in Matlab. Find an interactive, time-dependent comparison of the mutability map to the affinity map model here https://amitaiassaf.github.io/SpikeGeometry/SARSCoV2EvoT.html.

## Acknowledgments

This work was supported by the National Institutes of Health, grant number 2U19AI057229-16. The author thanks A. K. Chakraborty, D. Lingwood, A, Seeber, J. P. Barton, and K. Dao Duc for their helpful discussions and comments. The author thanks O. Gal for his contribution to the SARS-CoV-2 time spike evolution comparison tool.

## Code availability

The simulation and analysis code for this study are available at: https://amitaiassaf.github.io/SpikeGeometry/SARSCoV2EvoT.html.

## Supplementary information

### Supplementary Tables

**Table S 1.** Dimensions of the elements constructing the coarse-grained models. Description of the elements constructing the coarse-grained antibody model (S Figure 1C-D) and the immunogens (Figure 1A-i, B-i, Figure 3A-i, S Figure 1A,B).

| Index | Description | Size $\sigma_{i,i}$ | Bead color in Figure S1 |
| --- | --- | --- | --- |
| 1 | “Structural” atom of the immunogen. Not an epitope. The ferritin body is composed of such beads | 0.8nm | - |
| 2 | Fc of the Ab | 5nm | Blue |
| 3 | Hinge bead connecting the Fc bead to the two arms hinges | 1nm | Orange |
| 4 | Hinge beads in the Ab arms | 1.75nm | White |
| 5 | Fab | 1.75nm | Yellow |
| 6 | Arm bead in the Ab | 4.2nm | Magenta |
| 7 | Epitope beads | 1.75nm | - |

**Table S 2.** LJ interaction parameters. Values of *σ_i,j_* in nm.

| Bead index | 1 | 2 | 3 | 4 | 5 | 6 | 7 |
| --- | --- | --- | --- | --- | --- | --- | --- |
| 1 | - | 1.275 | 1.275 | 1.275 | 1.275 | 1.275 | 1.275 |
| 2 |  | - | 3.375 | 3.375 | 3.375 | 3.375 | 1.75 |
| 3 |  |  | - | 0.50 | 1.75 | 1.75 | 1.75 |
| 4 |  |  |  | - | 1.75 | 1.75 | 1.75 |
| 5 |  |  |  |  | - | 1.75 | 1.75 |
| 6 |  |  |  |  |  | - | 1.75 |
| 7 |  |  |  |  |  |  | - |

### Supplementary Figures

**S Figure 1.**
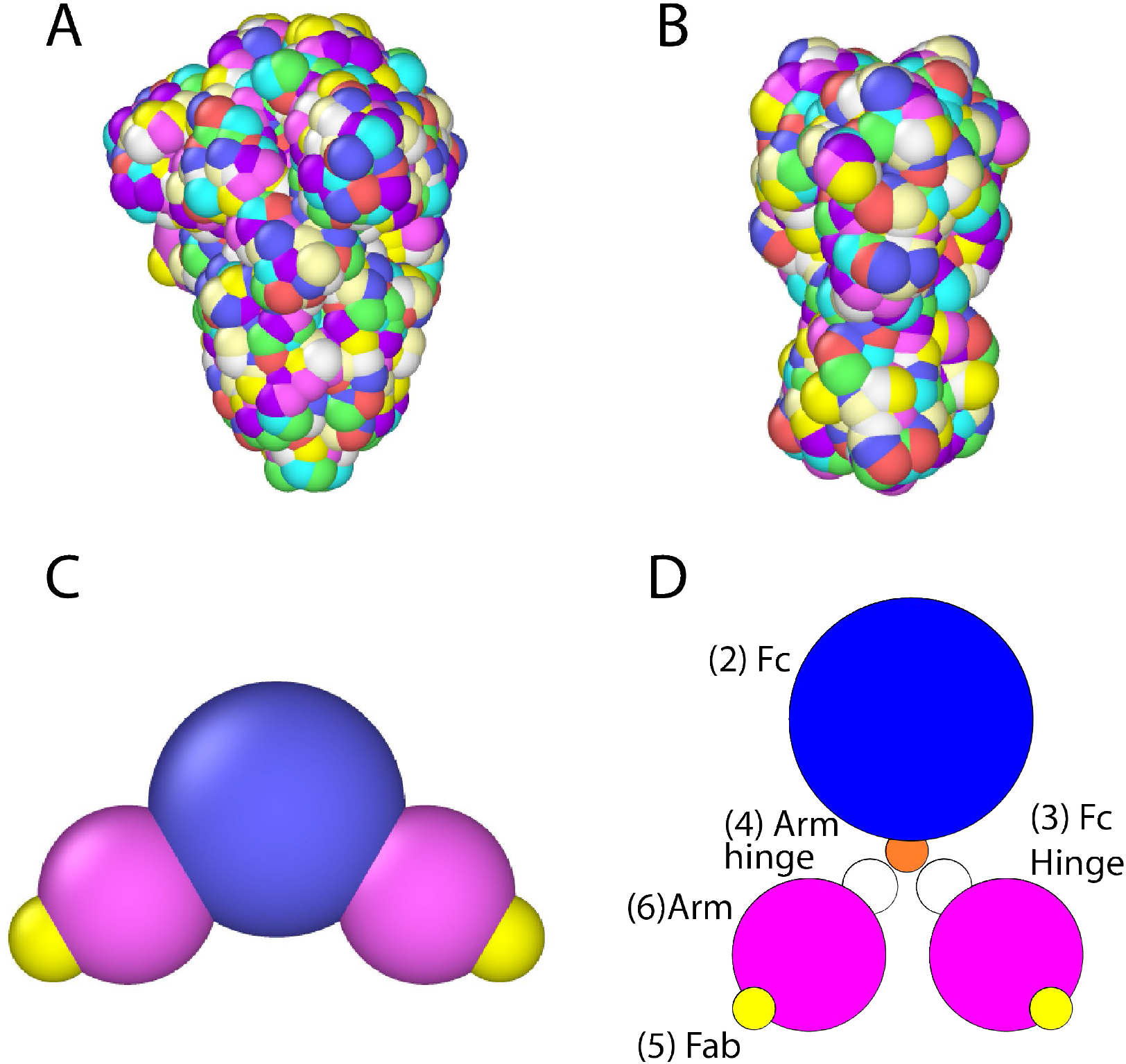
S and HA antigen-antibody models and antibody. **(A)** Set of 254 distinct residues on the surface of the S protein of SARS-CoV-2. See also Materials and Methods. **(B)** Set of 184 distinct residues on the surface of HA. **(C-D)** Schematic representation of the antibody (Ab). The large blue bead represents the Fc part of the Ab. The two magenta beads are the arms, and the yellow beads are the Fab section of the arms. The model also contains hinge beads between the Fc and the arms. For full description see “Coarse-grained model of the antibody”).

**S Figure 2.**
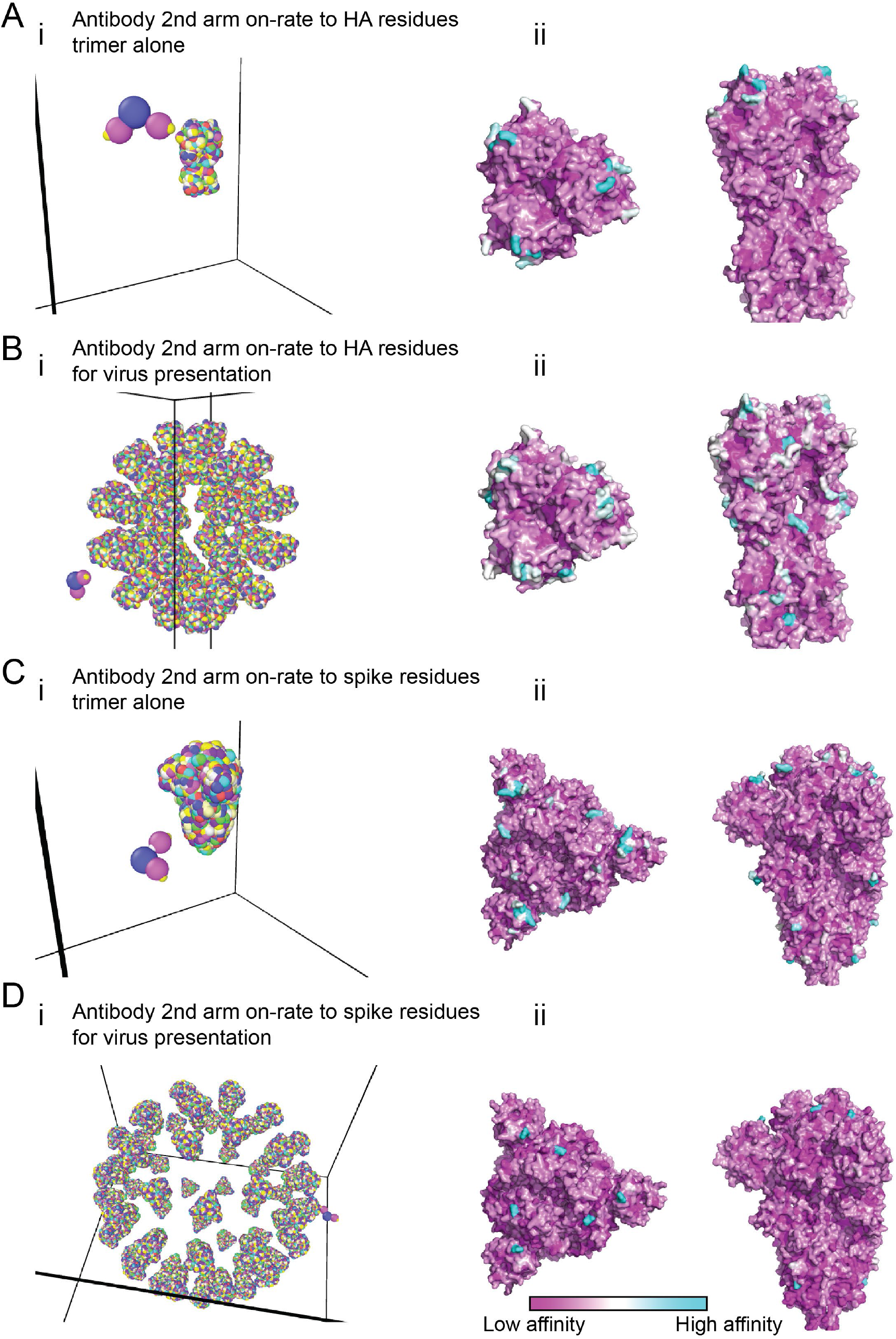
Second arm on-rate. Second arm binding was measured as on-rate to the second epitope, given that the first site is already bound (see eq.(S5)). **(A-B)** HA in its trimeric form (A) or on the surface of the virus (B) (see Figure 1A-B for details). (C-D) S protein in its trimeric form (C) or on the surface of the virus (D) (see Figure 3A for details). The second arm binding rate to cyan sites is high, intermediate to white sites, low for purple sites, and was the average over multiple simulations.

**S Figure 3.**
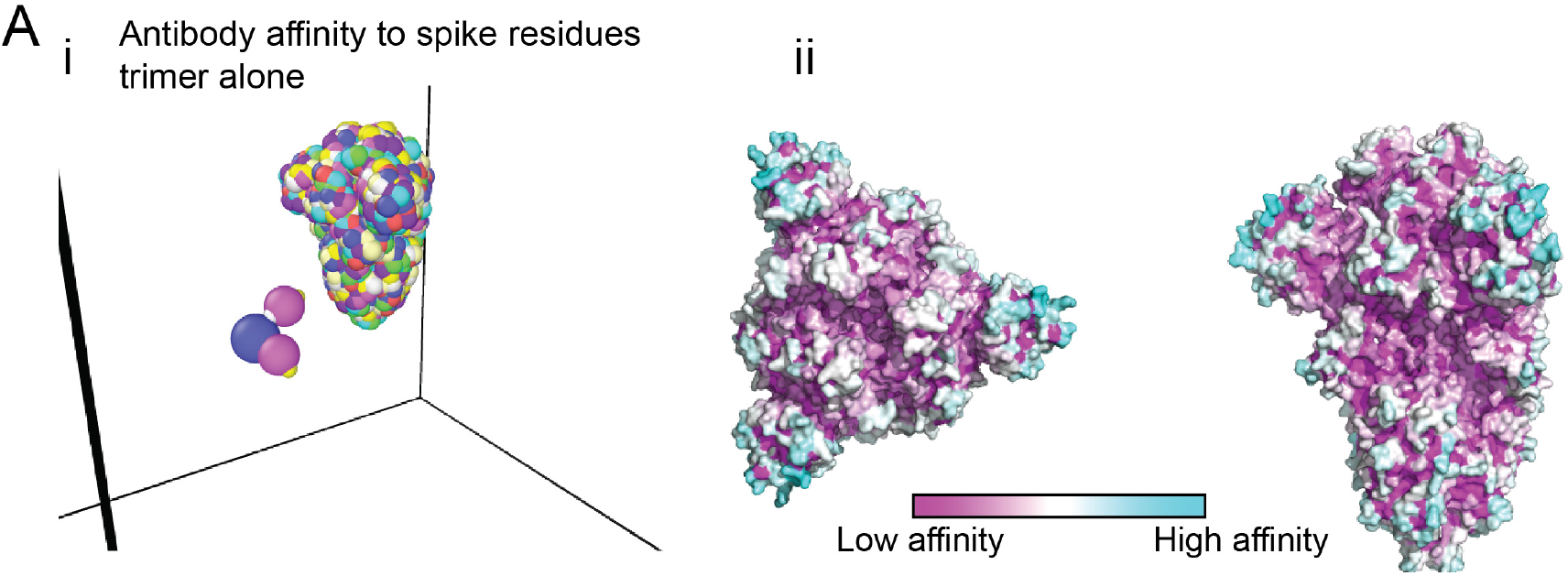
Antibody affinity to surface residues of the coronavirus spike in its trimeric form. **(A)** SARS CoV-2 spike (S) protein in its closed form (Walls et al., 2020). A detailed atomistic structure of the spike is coarse-grained and presented in rainbow colors (panel i). Every colored bead on the spike is a residue, representing a different S epitope (254 different possible sites on trimeric S). Panel ii depicts coarse-grained simulations for the Ab affinity to these residues (see Figure 1A-ii for definition and color-coding).

**S Figure 4.**
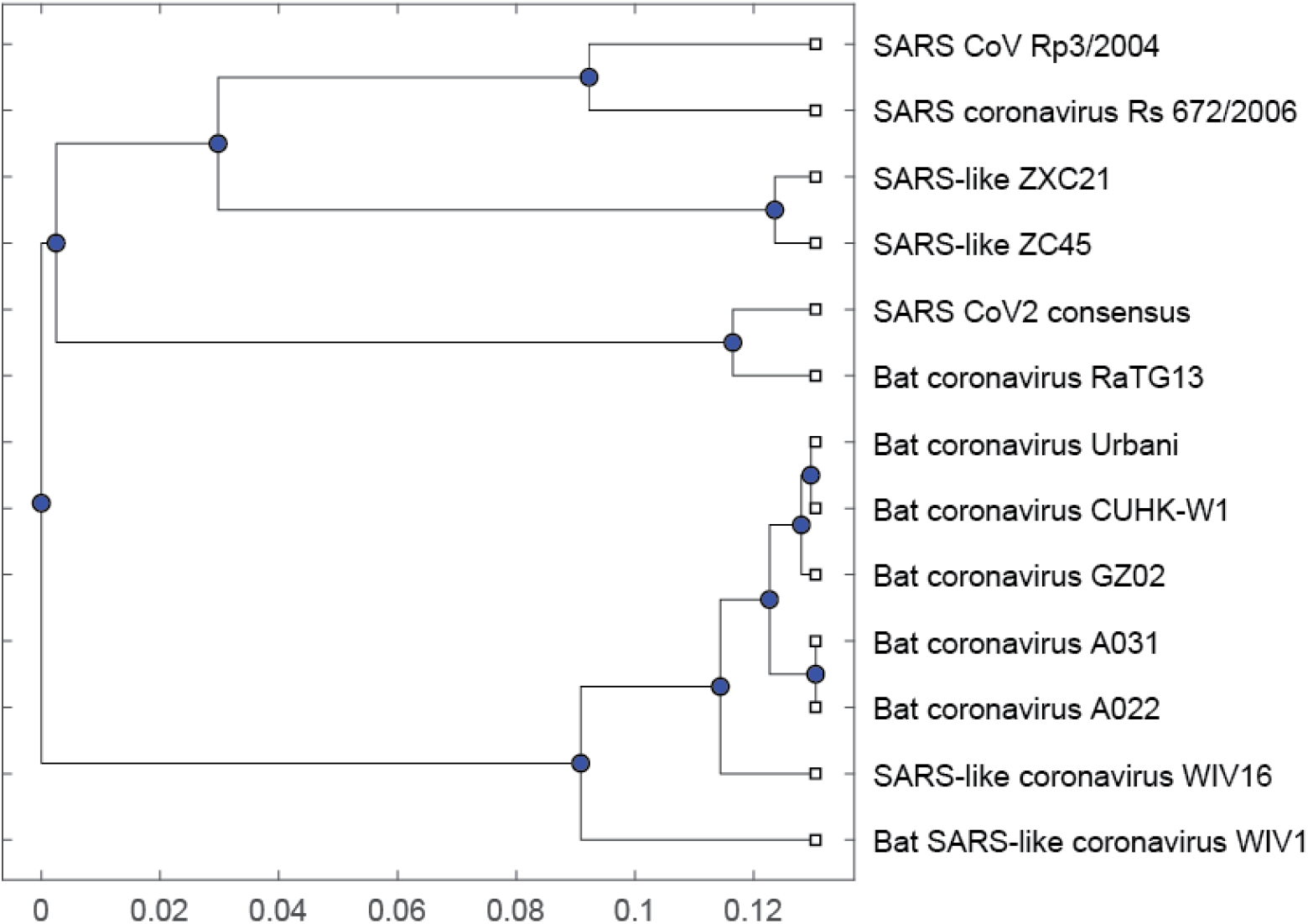
Sarbecovirus subgenus phylogenetic tree. The sequence origin is detailed in Table 1.

**S Figure 5.**
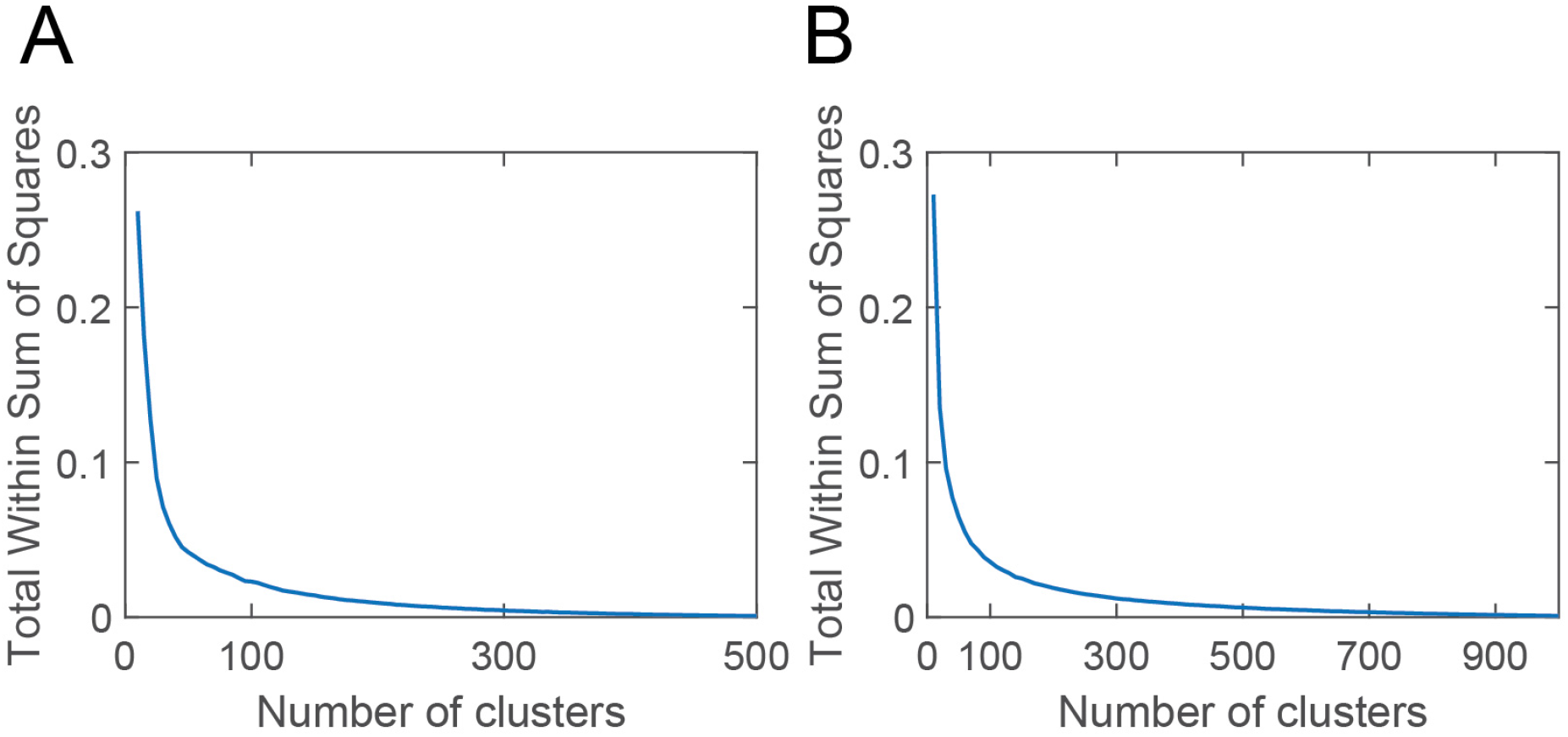
Clustering of spike surface residues (epitopes). **(A)** The Total Within Sum of Squares as a function of the number of clusters computed for seasonal influenza spike HA. Related to Figure 2 **(B)** The Total Within Sum of Squares as a function of the number of clusters computed for the sarbecovirus subgenus spike. Related to Figure 3.

